# Visual processing and attention rather than face and emotion processing play a distinct role in ASD: an EEG study

**DOI:** 10.1101/517664

**Authors:** Adyasha Tejaswi Khuntia, Rechu Divakar, Fabio Apicella, Filippo Muratori, Koel Das

## Abstract

Autism Spectrum Disorder results in deficit in social interaction, non-verbal communication and social reciprocity. Cognitive tasks pertaining to emotion processing are often preferred to distinguish the ASD children from the typically developing ones. We analysed the role of face and emotion processing in ASD and explored the feasibility of using EEG as a neural marker for detecting ASD. Subjects performed a visual perceptual task with face and nonface stimuli. Successful ASD detection was possible as early as 50 ms. post stimulus onset. Alpha and Beta oscillations seem to best identify autistic individuals. Multivariate pattern analysis and source localization studies points to the role of early visual processing and attention rather than emotion and face processing in detecting autism.

---

Autism Spectrum Disorder (ASD) is a developmental disorder characterized by learning difficulties. Children suffering from ASD show impairment in social communication, namely imitation, joint attention and empathy. Numerous studies have focused on understanding the neural mechanism leading to ASD (Hadjikhani et al. 2007; Rizzolatti and Fabbri-Destro 2010; Rubenstein and Merzenich 2003; Williams et al. 2001) but they still remain unclear. One hypothesis attributes deficit in motor processing resulting in a broken mirror neuron system (MNS) as the underlying cause of autism (Hadjikhani et al. 2007; Rizzolatti and Fabbri-Destro 2010; Williams et al. 2001). But the broken mirror system theory has come under criticism in recent times (Enticott et al. 2013). Several ASD studies have demonstrated atypical neural connectivity, both anatomical and functional (Anderson et al. 2011; Just et al. 2004, 2007; Villalobos et al. 2005), but the nature of the abnormality remains unclear.

Specifically, the role of face processing and emotion recognition in ASD remains a much-debated topic in the community (Campatelli et al. 2013). While many groups have found deficit in identifying different facial emotions to be one of the impairments in ASD (Arkush et al., n.d.; Bal et al. 2010; Buitelaar et al. 1999; Castelli 2005; Rump et al. 2009), there have been contradicting claims as well (Bookheimer et al. 2008; Ewing, Pellicano, and Rhodes 2013; Sterling et al. 2008; Tehrani-Doost et al. 2012). Literature involving neural studies of ASD and control population also report similar contradictions (Billeci et al. 2013). Typically, face processing deficit is related to an impairment in fusiform gyrus. Although several groups have shown less fusiform face area (FFA) activity during face processing (Corbett et al. 2009), others have negated such findings (Apicella et al. 2013; Hadjikhani et al. 2004). A specific evoked response potential (ERP) N170, has been established as a neural marker appearing during face detection (Anaki, Zion-Golumbic, and Bentin 2007) and/or processing (Barbeau et al. 2008; Halit et al. 2004). Similarly, another ERP component P1, is known to have a higher amplitude for a face stimuli (M. J. Herrmann et al. 2005; Martin J. Herrmann et al. 2005; Itier and Taylor 2004). Both N170 and P1 components have been used to study impairments in facial processing in autism studies (Batty et al. 2011; Hileman et al. 2011). These components, N170 and P1 were in fact shown to be similar for both the groups (Apicella et al. 2013; Webb et al. 2010). Parallel opposing studies claiming a higher latency for neurotypical subjects in similar facial ERP components have also been addressed (McPartland et al. 2004; K. Pierce et al. 2001). The P1 component is interestingly known to be affected by individual’s attentive state during task performance (Mangun 1995). Additionally, task load requiring higher attention (Taylor 2002) and selective attention (Luck et al. 1994) have also been reported to affect P1. The role of the evoked P1 component in autistic individuals thus remains inconclusive; whether it is arising because of a deficit in face processing or as a result of the different attentional requirements while observing the stimuli. Recent findings investigating the role of P1 and N100, a component associated with attentional orientation (Heinze et al. 1990), have shown a slower P1 latency and weaker N100 amplitude among ASD individuals (Wang et al. 2017), thereby, indicating abnormalities in attention modalities for the non-typical group. But, studies on P300 activity, an important component contributing to sustained attention to target, alertness and attention allocation during tasks, attribute a similar P300 activity for both ASD and control groups (Gomot and Wicker 2012). Several other eye-tracking based experimental studies have addressed abnormalities among ASD individuals pertaining to visual-attention based tasks (van der Geest et al. 2001; Guillon et al. 2014; Townsend, Harris, and Courchesne 1996).

Other neuroimaging, ERP and eye-tracking literatures still fail to explicitly resolve whether ASD individuals have a reduced efficacy in encoding facial structures or whether the observed differences in neural signals are an outcome of disparity in the extent of visual processing pertaining to how efficiently the respective individuals are processing what they see. These findings have been partially beneficial in exploring the role of emotion processing in autistic subjects. Notably, previous research have reported ASD individuals to have a higher tendency in recognizing familiar faces in contrast to a lower performance for strangers with their performance being at par with the control group (Bal et al. 2010; Karen Pierce and Redcay 2008). In another study, autistic children were required to recognize six types of emotions. These subjects were able to recognize the emotions, although with different intensity levels and showed similar errors as control (Buitelaar et al. 1999; Castelli 2005). Hence the exact extent of facial emotion processing in ASD remains unclear.

In the current study, we have focussed on the role of face perception and emotion processing in children diagnosed with ASD. Using a visual perceptual task with faces and non-faces (Apicella et al. 2013), we performed a systematic analysis to explore the early effect of both face and emotion processing in ASD and control population. Our study suggests the role of visual processing and attention in successfully detecting ASD. Diagnosis of ASD is typically carried out by employing behavioural markers but use of reliable neural markers could potentially result in immense advancement in the field of ASD detection. Here, we have also tried to explore the possibility of finding reliable neural markers for ASD with single trial EEG analysis using multivariate pattern analysis (MVPA). We have analysed neural data in both time and frequency domain and identified specific spatio-temporal regions which can potentially act as neural markers to identify subjects affected by ASD. Interestingly, beta oscillations along with alpha waves seem to best capture the distinct characteristic features of ASD, sometimes even before the stimulus onset. Source reconstruction have been used to explore the localized brain regions that potentially codes the difference between autistic and neurotypical subjects.

## 1. Methods

### 1.1. Stimuli and Display

All the participants were presented with 3 kinds of stimuli including faces, tree and cartoon. The face stimuli consisted of 3 kinds of emotional expressions: happy, fear and neutral and was obtained from a widely used database (Tottenham et al. 2009). 10 faces belonging to 10 subjects (5 male and 5 female) for each emotion was selected from the database. In order to avoid biased focus on external facial features, the features (ear, neck and hair) were deleted before presenting to the subjects. Even the tree stimuli were cropped to make them all of similar size. The face and tree stimuli were all converted to greyscale to avoid a bias. But, for cartoon stimuli, the colour was kept for retaining the subject’s attention on the screen. Tree was chosen as a non-face stimulus because of its similar attributes to face in regards of similar genetic variability, multidimensional and environment factors.

### 1.2 Observer and Experiment

The current study explored the high-density EEG data of 24 subjects. 12 autistic and 12 neurotypically developing age-matched individuals performed the experiment. The diagnosis of ASD within the participating group followed the DSM-5 criteria confirmed by ADOS-2 and ADI-R.

These children ranged within 6-13 years of age with a mean age of 10.2 years. The control group had an average age of 9.7 years. The subjects selected were maintained to be of similar intellectual level (total IQ≥80, VCI and PRI ≥ 70 in WISC-IV), absence of any known epileptic condition and genetic disorder. Written consents were obtained from all parents following due procedure.

The experimental paradigm consisted of 4 blocks lasting for 5 minutes in total for each individual with 3 pauses in between. Each of the blocks consisted of 100 images including 30 face stimuli (10 happy, 10 fearful and 10 neutral), 10 tree stimuli and 10 cartoon stimuli with each repeating twice. The task for the participants was to press a button on observing a cartoon stimulus on the screen with their preferred hand. Cartoon stimuli required explicit response from the children and reward was given based on their performance. Neural signals corresponding to cartoon stimuli were excluded from further analysis. Thus, the experimental paradigm rules out the unnecessary task-related effects like inattention, configuring a purely implicit, covert facial emotion perception task. Fig 1 illustrates the experimental paradigm followed to acquire the data. The stimuli were presented for 850 ms each in a random order and interleaved with a fixation cross with a randomized duration of 500–1500 ms to avoid expectation effects. The subjects were asked to look at a fixation cross appearing at the centre and in each corner of the display before the task commenced to have a reliable idea of gaze target when reviewing data. During the task administration, subjects were frequently asked to look at the fixation cross when it appeared. This was a strategy to induce them to focus on the incoming stimuli. In order to track where the subjects were looking during task performance, a firewire digital camera was held above the screen, positioned so as to capture individualistic performances.

**Fig 1.**
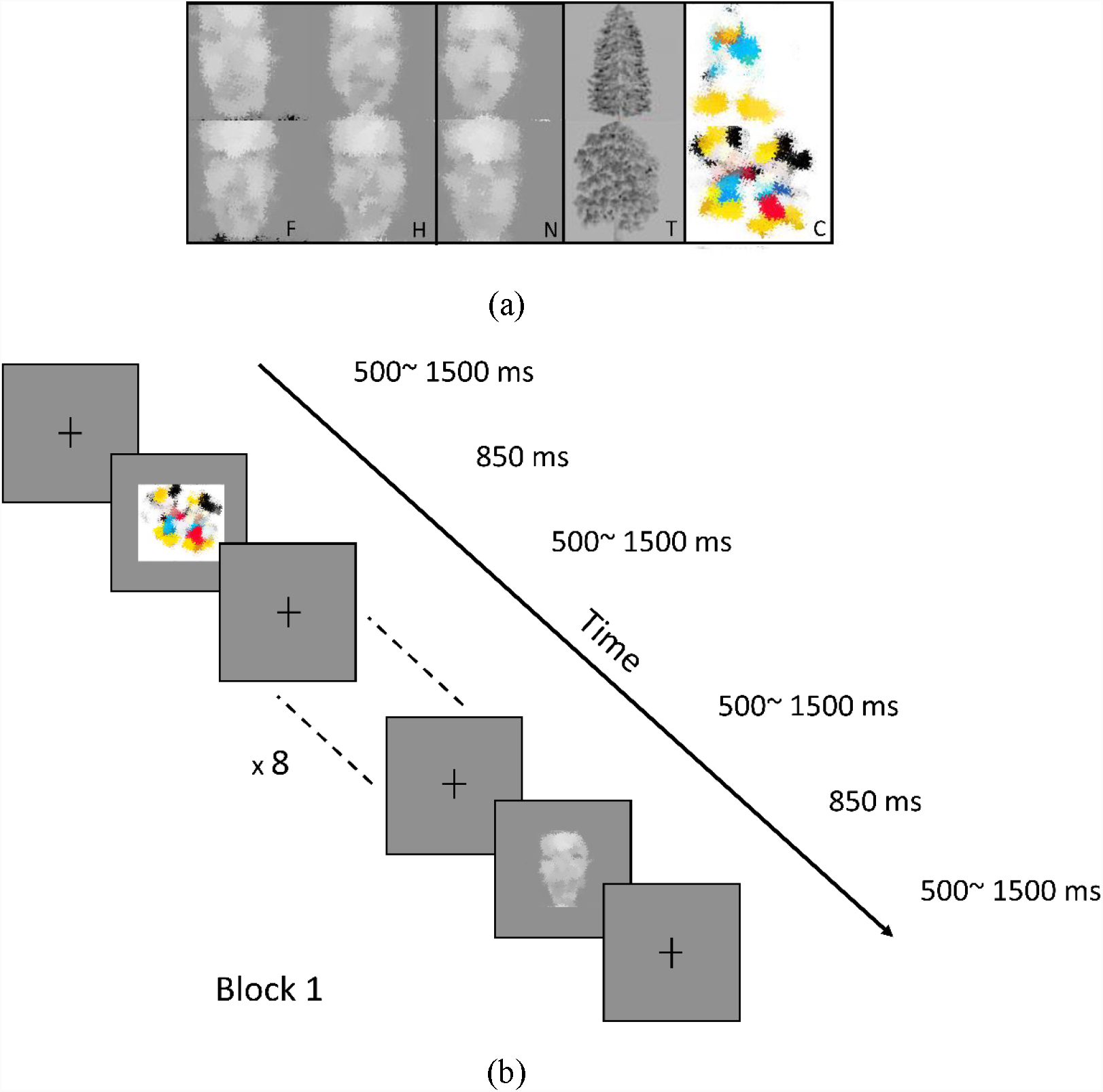
a) Stimuli-Emotional face: ‘fear’, ‘happy’; Non-emotional face: ‘neutral’; Non-face: ‘tree’. The images are obtained from a widely used database (Tottenham et al. 2009) b) Experimental Paradigm (The images are were obscured as per bioRiv policy)

### 1.3 Data Processing and Analysis

#### EEG Data Acquisition and Preprocessing

The data was acquired using a 128 – channel Hydrocel Geodesic Sensor sampled at 250 Hz. Impedances were maintained below 50 kΩ. Neural signals 150 ms prior to stimulus onset were retained as pre-stimulus signals. The data was bandpass filtered between 0.1 and 40 Hz using 3rd order zero phase Butterworth filter and referenced using average referencing. Trials containing ocular artefacts like eye movements and blinks were excluded. These were detected by analysing the bipolar electro-oculogram (EOG) with amplitudes exceeding +/- 100 mV or by visual inspection. Average time-locked ERPs were computed for different stimuli including 150 ms pre-stimulus baseline and 850 ms post-stimulus interval.

#### EEG Analysis

Data analysis was carried out using both classic event related potential (ERP) measures and multivariate pattern classifier. ERP was extracted from the baseline removed pre-processed data by averaging all the trials for each of the two groups, ASD and Typically Developing (TD) across each stimulus condition.

Multivariate pattern analysis was carried out by using Classwise Principle Component Analysis, CPCA, (K. Das, Osechinskiy, and Nenadic 2007) because of its superior performance in classifying high dimensional noisy neural signals with fewer training samples (K. Das, Osechinskiy, and Nenadic 2007; King et al. 2015; McCrimmon et al. 2017). Similar results were also obtained using linear support vector machine (SVM) but CPCA was used because of its faster execution time. CPCA is a two-step technique where the first step consists of identifying and discarding a sparse non-informative subspace in the data. The second step follows a linear combination of the remaining data in the chosen subspace to extract the best features suited for the efficacy of the classifier performance in the remaining subspace. 10-fold stratified cross-validation (Kohavi 1995) was used to report the classifier performance. The classifier was trained using 90% of the data while the remaining 10% was used for testing. The process was repeated 10 times to cover the entire span of data and the average classifier performance was reported. Classification performance was reported by calculating Area under the Receiver Operating Curve (AUC) (Mason and Graham 2002). The error bars for the classifier performance for each stimulus was calculated from the standard error across the AUC for 10-fold cross validation. Analysis was carried out in both time-domain and time-frequency domain.

#### Time Domain Analysis

The pattern classifier discriminated between ASD and TD subjects based on single trial multivariate discriminant analysis. The entire EEG data consisting of 150 ms pre-stimulus and 850 ms post-stimulus for all channels were used as input to the classifier which then discriminated between the two classes to give us a measure of its performance in terms of AUC.

Additionally, the data set was divided into 25 temporal windows consisting of 40 ms each to accurately predict the time windows with the highest classification. The classification rates were computed for each window to elucidate the temporal epoch that codes the difference between ASD and TD subjects when viewing different types of stimuli.

#### Time-frequency Domain Analysis

The acquired neural data for all the stimuli were transformed into the respective time-frequency domain using Short-term Fourier Transform to gain an insight on the behaviour of different frequency bands. Spectrograms were plotted to visualise the time-frequency plots for each channel for each time point for all the subjects. Time-frequency data was extracted for the different frequency bands: delta, theta, alpha, beta and gamma. Similar classifications (as time domain) were carried in the time-frequency domain using CPCA. The time points were divided into 23 windows and classification was carried out in each of them.

#### Discriminative Activity

Feature extraction matrices of CPCA can be viewed as a spatio temporal filter that produces the discriminative activity for each channel and time points (Koel Das, Rizzuto, and Nenadic 2009). Scalp topographic plots were used to visualize the spatio-temporal discriminatory regions. The multivariate pattern classifier CPCA only selects the most informative subspace while discarding the other. The filter coefficients for such a subspace with maximum features are informative ways to elucidate the spatio-temporal information regarding the locations which encode the distinguishing features of the two classes for each stimulus. Such topographic representations have been constructed in both time and time-frequency domain.

#### Source Localisation

To obtain the distinct localised regions in the brain for distinguishing the non-typical from the typical individuals, Source Reconstruction was performed using a Multiple Sparse Prior algorithm (Friston et al. 2008) implemented in SPM12 (http://www.fil.ion.ucl.ac.uk/spm/). The standard Montreal Neurological Institute (MNI) template was used as head model and the sources were identified using the automated anatomical labelling (AAL) atlas. The difference wave was computed by subtracting the ERP for ASD from that of TD. The source activity was estimated by giving the difference wave for each stimulus as input and the resultant source activity maps were converted into 3D images. The source was estimated in the peak discriminating time range construed from the single trial EEG analysis. The estimated source regions for each type of stimuli were analysed systematically.

## 2. Results

### 2.1. Univariate Analysis

Grand averaged Event Related Potentials are used to visualize any differences arising from the different classes of stimuli for ASD and control group. Waveforms for electrodes in the frontal (FC6, F4, F6, FT8, AF4), central (C4, C2) and occipital (OZ, POZ, O1, O2) areas have been shown in Fig 2 for an emotional face (fear) and a non-face stimuli (tree). As evident from the figure, there is a difference between the two groups as early as 100 ms post stimulus onset for both tree and fearful face. Similar ERP response are observed for all other stimuli as well. The difference in response between the two groups are sustained from 100 ms onwards and there is no specific difference in the N170 component for face stimuli for the ASD group. The ERP results indicate that perhaps early face processing mechanism is not impaired in case of subjects suffering from ASD.

**Fig 2.**
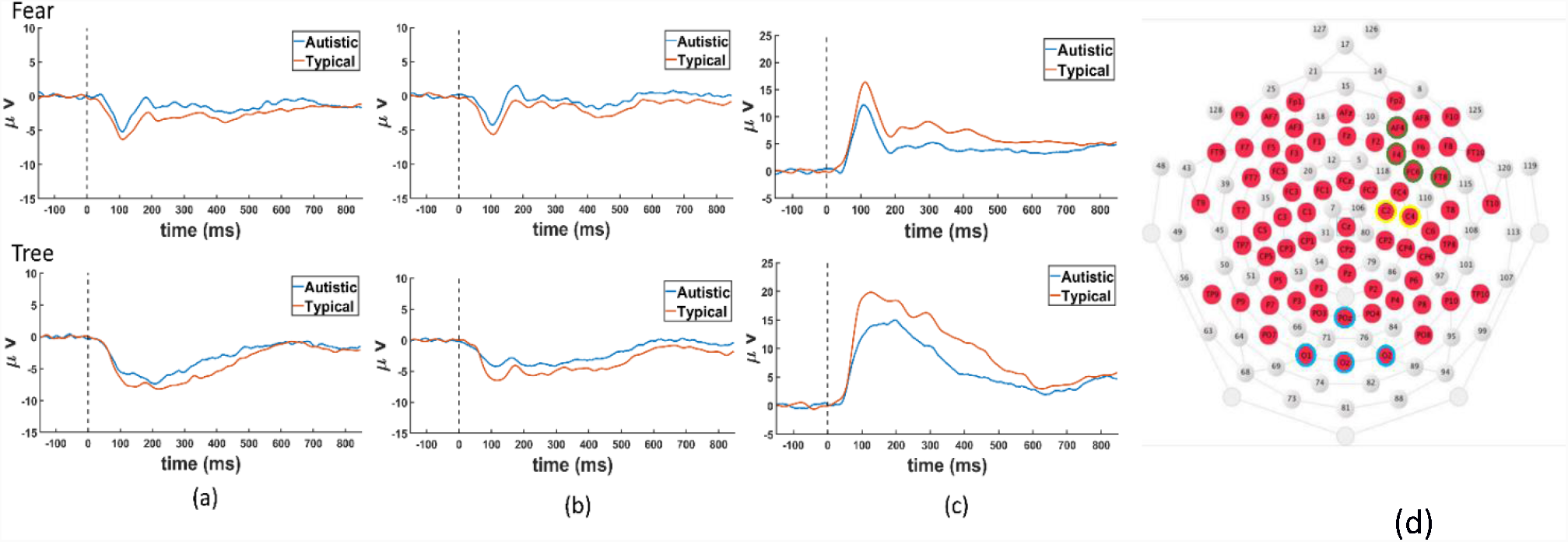
Event related potentials corresponding to Fear and Tree stimuli of electrodes (a) FC6, F4, F6, FT8, AF4 (b) C4, C2 (c) POz, Oz, O1, O2 (d) Channel location of 128-channel HGSN EEG cap used in the experiment. The channel marked in green, yellow and blue correspond to the electrodes mentioned in (a), (b) and (c)

### 2.2. Multivariate Pattern Analysis

#### Time Domain

We used CPCA to classify the EEG data into ASD and TD for different types of stimuli (Fig 3). A 10-fold stratified cross validation was performed on the single trial EEG data for different stimuli. The high classification accuracies (76-80%) of the entire time-series data for all the conditions demonstrates the viability of using single trial EEG as a potential neural marker for ASD detection.

**Fig 3.**
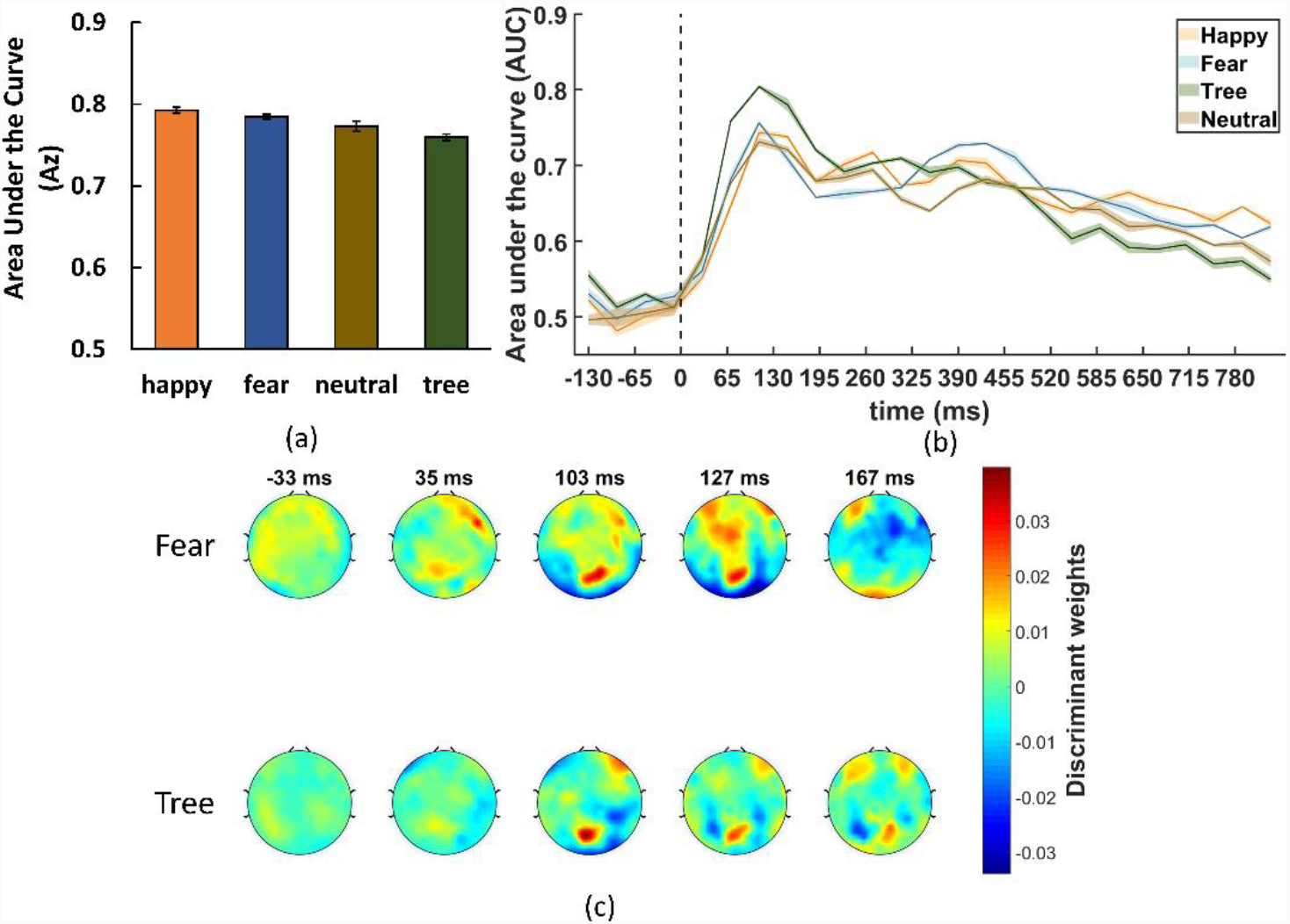
(a) Area under the curve using pattern classifier for all stimuli in the time domain (b) AUC in 40 ms time windows for all stimuli to infer the time interval with the maximum discriminant activity (c) Scalp topographic plots as a measure of the discriminate spatio-temporal regions for fear and tree stimuli. The color intensity encodes the discriminant weights or filter coefficients to elucidate spatio-temporal discriminant features between the two groups. Discriminant activity is greater in areas having highly positive or highly negative values. The discriminant weights are shown in the colorbar. All plots are generated using the EEGLAB (Delorme and Makeig 2004) using all the electrodes

Next, we analysed the EEG response by breaking the signal into 40 ms time windows and use CPCA for classification in each time window. A higher classification rate at an earlier epoch might suggest that less complex cognitive processes involving attention and visual information are responsible for the observed differences in activity among the two groups. But typically difference distinguishing the two groups due to factors including emotion processing and face processing occurs at a later time window (Dawson, Webb, and McPartland 2005; Lerner, McPartland, and Morris 2013). Highest classification rate was observed in the time interval of 50-130 ms for all the conditions. Classification rates are significantly greater than chance even after 130 ms and even for the pre-stimulus for almost all stimulus condition (p<0.05, FDR corrected). In order to explore contribution of the spatio-temporal areas that play a role in discriminating between the two classes, the discriminant activity has been represented as topoplots in Fig 3c. The discriminatory activity (shown as highly positive or highly negative values) initially appears in the occipital lobe and post 100 ms moves towards frontal lobe for all the conditions. Interestingly, no significant differences between emotional and non-emotional stimuli are observed (F (3,96) =0.21, p=0.888).

#### Time-frequency domain

Further analysis of the neural data was carried out in the time-frequency domain to analyse the effect of different frequency bands (Fig 4). Spectrogram analysis showed a higher difference in alpha and beta activity while results did not vary for face and non-face stimuli. Fig 5 shows classification results for all the stimulus and all the bands. Classification was highest for alpha and beta waves. The accuracy of the classification ranges from 79-84% for alpha wave and 77-90% for beta wave. No discernible difference was observed in the classification trend across the stimuli. Time window analysis gave high classification performance for all frequency bands in all windows. Interestingly, pre-stimulus signals also produced classification accuracy of around 70% for all conditions. Fig 4 (b) illustrates the classification trends for alpha frequency band for all the conditions.

**Fig 4.**
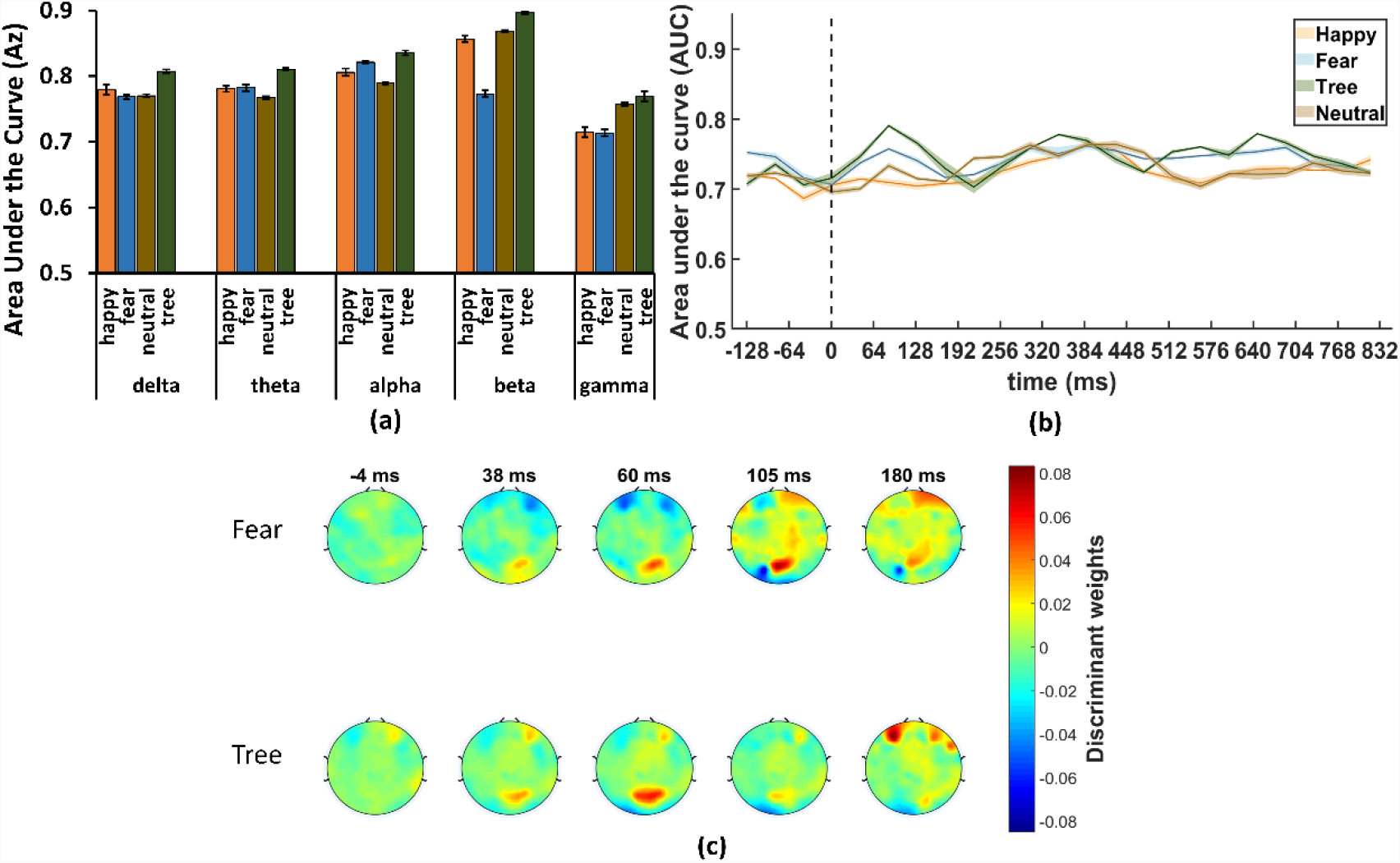
(a) Area under the curve for the performance of the classifier for all stimuli in the time-frequency domain (b) AUC across 23-time windows for alpha band (c) Scalp topographic plots as a measure of the discriminate spatio-temporal regions for alpha band of fear and tree stimuli generated similar to the topographic maps in time-domain

**Fig 5.**
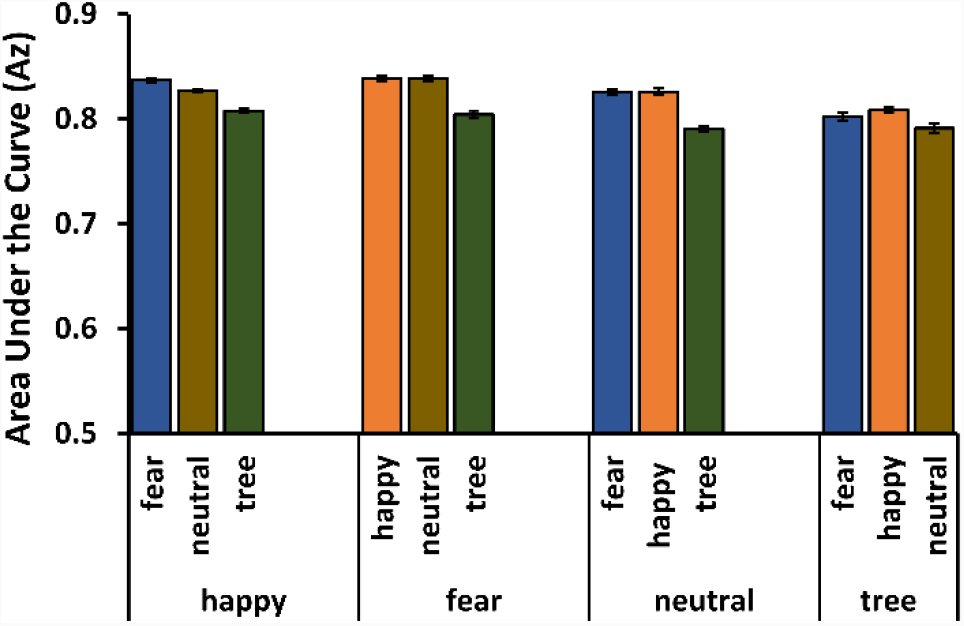
Area under the curve for the classifier when it was trained on one stimulus and tested on all other stimuli. High classifier accuracy points to insignificant role of stimuli type in ASD detection

The spatial representation of the discriminant activity (Fig 4 (c)) shows similar trends as the time domain data for both emotional (fear) and non-emotional (tree) stimuli occurring at occipital and frontal lobe. There was no systematic difference in MVPA performance or localised discriminability between the face and non-face stimuli.

In an additional study to analyse the effect of face/non-face, emotional/neutral stimuli, we systematically trained our pattern classifier on one type of stimuli, and tested with another type. The results of the analysis are given in Fig 5. From the results, it follows that ASD and TD groups can be classified with very high accuracy even when the classifier is trained on a different stimulus. MVPA performance for all cases are around 81% in the time domain and 84% in the frequency domain. The above results seem to point to the absence of face and emotion processing in discriminating between the ASD and TD subjects.

### 2.3. Source Localisation

Finally, we performed source localisation (SL) or source reconstruction to obtain the regions in the brain which result in the observed evoked response. Since, the classification rates using EEG signals were highest in the range 50-130 ms, the active spatial areas for this specific time window was localised using the difference ERP wave. The areas showing activity for most of the stimuli were Lateral Orbital Gyrus (LOrG), Inferior temporal gyrus (ITG) and Inferior Occipital Gyrus (IOG). Source localization results for emotional face and non-face stimuli are provided in Fig 6. The source analysis revealed that similar areas contributed to difference between the two groups irrespective of stimuli provided and this finding is consistent with our previous results (Figs 3c, 4c). The source areas specifically LOrG have been previously implicated in attentional processing (Hartikainen, Ogawa, and Knight 2012) and reconstructed sources point to deficit in visual and attentional processing as the main difference between ASD and TD subjects.

**Fig 6.**
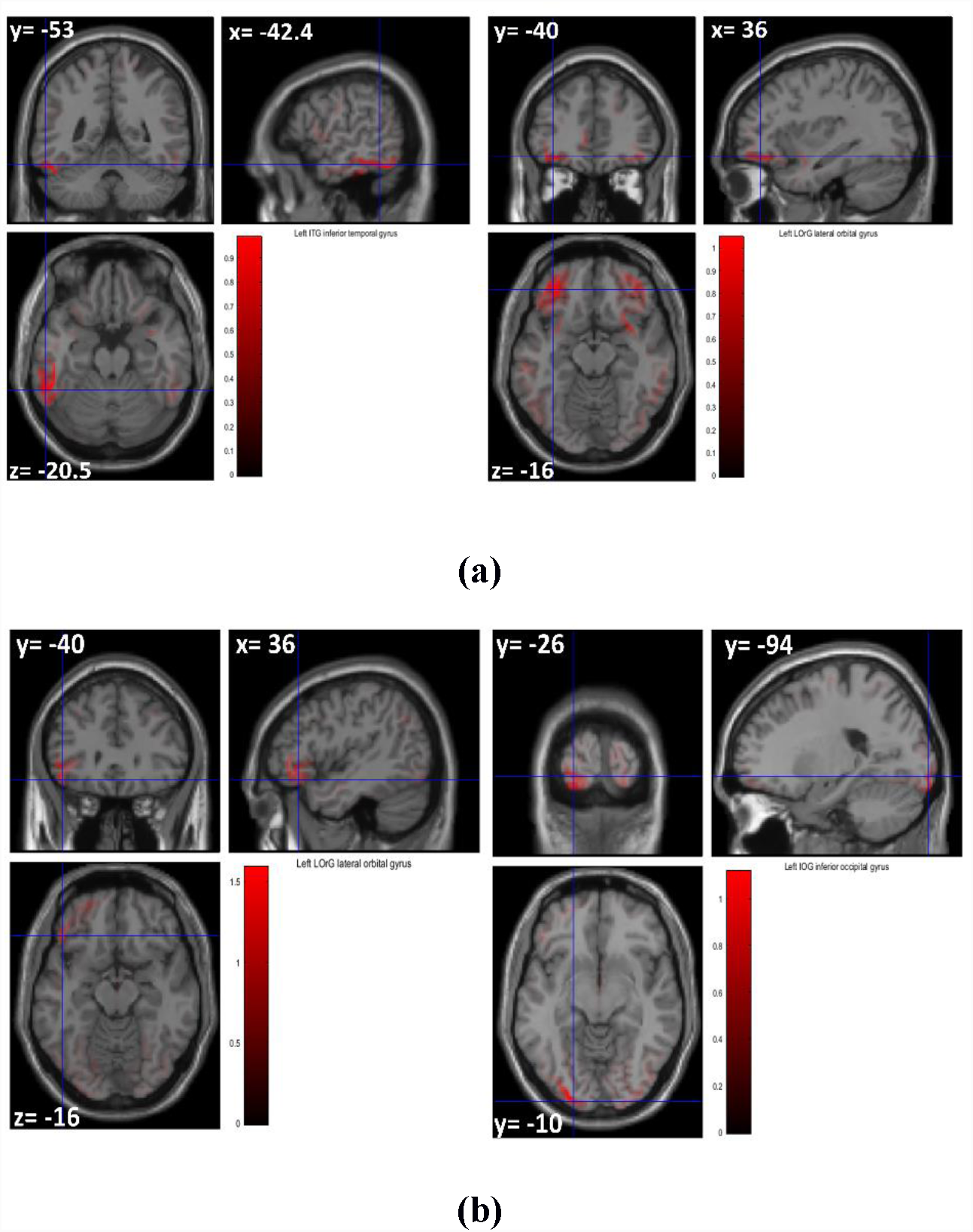
Source Reconstruction of the difference ERP for the significant time window of 50-130ms for (a) Fear (b) Tree stimuli. Significant source area includes Lateral orbital gyrus (LOrG) and Inferior temporal gyrus (ITG) activity for ‘Fear’ shown in (a) and LOrG, Inferior occipital gyrus (IOG) for ‘Tree’ shown in (b). The slice co-ordinates mentioned are in mm’s in MNI space

## 3. Discussion

### 3.1. Neural marker for ASD

Autism diagnosis typically takes place during pre-school years but it is known that manifestation of disorder starts much earlier (Oosterling et al. 2010). However, absence of reliable markers for ASD poses a challenge and in this study, we explore the feasibility of finding a neural marker for autism detection. From the results, we can reliably demonstrate that it is possible to detect ASD with high accuracy using only single trial EEG. The results furthermore demonstrate that it is possible to detect ASD and neurotypical children using only first 50-130 ms post stimulus onset EEG signal. Our additional analysis (Fig. 5) suggests that irrespective of the stimulus, single trial EEG detects subjects suffering from ASD with high accuracy (>80%). The results presented here alludes to the possibility of developing an experimental paradigm wherein rapid detection of ASD and neurotypical subjects using less than 200 ms of EEG recording is possible.

### 3.2. Face and Emotion Processing in ASD

The role of face processing and emotion processing has been a contentious issue within autism research community (Arkush et al., n.d.; Bal et al. 2010; Bookheimer et al. 2008; Castelli 2005; Ewing, Pellicano, and Rhodes 2013; Sterling et al. 2008; Tehrani-Doost et al. 2012). Eye-tracking studies relating to emotion regulation in ASD highlight an atypical response on seeing faces of others by the ASD subjects (Arkush et al., n.d.; Bal et al. 2010). It has been reported that children with autism could recognise all six emotions although with different intensity levels and also followed similar errors as control (Buitelaar et al. 1999; Castelli 2005). Again, a study by Tehrani et al. showed that youth with ASD have similar ability to recognise faces (Tehrani-Doost et al. 2012). Underlying attention for face processing of familiar and non-familiar faces by eye tracking showed a similar accuracy for both the control and ASD groups (Sterling et al. 2008). In the current study we wanted to focus on the neural mediators discriminating ASD from neurotypical subjects using face/non-face stimuli. We explored the role of face processing and emotion processing in ASD and neurotypical subjects and found no significant difference in face and emotion processing between subjects belonging to both groups. Our result is consistent with the findings using same stimuli (Apicella et al. 2013) wherein the authors could not find any significant differences in early face-sensitive ERP components between ASD and neurotypical children. Our results further show even a non-face stimulus like tree gave similar superior performance while discriminating between the two groups. We found face sensitive ERP components like N170 to be similar for both the groups which is consistent with previous studies (Apicella et al. 2013; Webb et al. 2010). There were no significant differences in the latencies of P1, N170 for all stimulus conditions.

We demonstrated that the multivariate pattern classifier was able to detect ASD subjects with high accuracy even when the classifier was trained on one type of stimulus and tested with others. MVPA results seem to insinuate the insignificant role of stimuli in detecting ASD and suggest that deficit in visual processing rather than face and emotion processing play a key role in ASD detection.

### 3.3. Role of attention in ASD

ASD individuals have been known to have abnormalities in visual attention modalities. They have slower P100 latencies, weaker N100 amplitude and larger P300 amplitude during exposure to visual stimuli (Wang et al. 2017). But there have been contradictory claims showing a similar P300 activity for both ASD and control population (Courchesne et al. 1985; Pritchard, Raz, and August 1987). ASD individuals have been reported to evoke similar P3b latencies but with less amplitude (Gomot and Wicker 2012). ASD individuals have also been known to overprocess information for differentiating target and novel stimuli successfully (Sokhadze et al. 2009). These findings elucidate an attentional abnormality for ASD children while performing task-based experiment. Our results also seem to suggest the role of visual attention as the primary discriminatory factor between ASD and neurotypical subjects. Our results show a higher classification accuracy for the beta waves which are known to play a role in attention (Siegel et al. 2008; Vázquez Marrufo et al. 2001). An early prediction before 200 ms also supports the role of attention in discriminating between subjects. Alpha wave activity is known to differ for the two groups (Dickinson et al. 2018; Keehn et al. 2017) and our classification analysis using alpha waves corroborates the same. Interestingly we also found high classification accuracy even in pre-stimulus signals especially in time-frequency domain suggesting that type of stimulus may not play an important role in detecting ASD using neural data. The source localization results show that lateral orbital gyrus (LOrG), is localised for all types of stimuli. This region has been reported to play a role in attention processing (Hartikainen, Ogawa, and Knight 2012). We also found some activity in inferior temporal gyrus (ITG) for all face stimuli and in inferior occipital gyrus (IOG) for fear, happy and tree stimuli. IOG have been previously reported to contribute to visual processing (Fockert et al. 2001). While ITG primarily processes face and objects, earlier results showing a heightened activation of ITG has been reported (Schultz et al. 2000) possibly due to a reduced attention to faces (Dawson et al. 2002). It is interesting to note that face fusiform area (FFA) did not show up as the prominent discriminatory source for most of the face stimuli which suggests that face processing ability might not play a key role in differentiating between ASD and our source localization is consistent with the pattern classifier results and possibly alludes to the role of visual attention, rather than face and emotion processing, in coding the difference between autistic and typical subjects.

## 5. Conclusion

Our current study suggests the possibility of using EEG signals as a potential neural marker for detecting ASD and neurotypical subjects. Our findings imply a deficit in attention and visual processing instead of the more complex, emotion processing and face processing in subjects suffering from ASD. It is possible to detect ASD subjects using a visual perceptual task irrespective of the type of face/non-face stimuli. The localised regions obtained by source reconstruction analysis suggest that attention and visual processing play a crucial role in differentiating between ASD and neurotypical subjects. Future studies might be devoted to initially designing an experimental paradigm with larger group of subjects for developing a neural marker for autism. Second, they could be tested in younger population: in fact autistic symptoms develop as early as the first year (Oosterling et al. 2010). Nevertheless, even with the advent of early screening (Daniels et al. 2014), there still remains a delay between ASD manifestation and diagnosis; this delay leads to failure to provide early intervention and improved results. Under the circumstances, use of early neural markers for ASD can potentially result in earlier intervention thereby resulting in better prognosis.

### Ethical approval

All procedures performed in studies involving human participants were in accordance with the ethical standards of the institutional and/or national research committee and with the 1964 Helsinki declaration and its later amendments or comparable ethical standards.

